## Supplemental File 1 for "Molecular function limits divergent protein evolution on planetary timescales"

**Supplementary file 1.** Considered model species and pairwise average divergence times ^a,b,c,d^ (in billions of years)[17].

|  | Eukarya | | | | | | | | | | Bacteria | | | | | | | | | Archaea | | |
| --- | --- | --- | --- | --- | --- | --- | --- | --- | --- | --- | --- | --- | --- | --- | --- | --- | --- | --- | --- | --- | --- | --- |
|  | *Homo sapiens* | *Rattus norvegicus* | *Xenopus laevis* | *Danio rerio* | *Gallus gallus* | *Drosophila melanogaster* | *Saccharomyces cerevisiae* | *Caenorhabditis elegans* | *Arabidopsis thaliana* | *Oryza sativa* | *Bacillus subtilis* | *Thermotoga maritima* | *Pseudomonas aeruginosa* | *Mycobacterium tuberculosis* | *Yersinia pestis* | *Haemophilus influenzae* | *Salmonella enterica* | *Escherichia coli* | *Agrobacterium fabrum* | *Pyrococcus horikoshii* | *Sulfolobus solfataricus* | *Haloferax volcanii* |
| *H.sapiens* | 0.0 | 0.02 | 0.01 | 0.03 | 0.01 | 0.23 | 0.24 | 0.23 | 0.17 | 0.17 | -- | -- | -- | -- | -- | -- | -- | -- | -- | 0.05 | -- | 0.05 |
| *R. norvegicus* | 0.1 | 0.0 | 0.01 | 0.03 | 0.01 | 0.23 | 0.24 | 0.23 | 0.17 | 0.17 | -- | -- | -- | -- | -- | -- | -- | -- | -- | 0.05 | -- | 0.05 |
| *X. laevis* | 0.4 | 0.4 | 0.0 | 0.03 | 0.01 | 0.23 | 0.24 | 0.23 | 0.17 | 0.17 | -- | -- | -- | -- | -- | -- | -- | -- | -- | 0.05 | -- | 0.05 |
| *D. rerio* | 0.4 | 0.4 | 0.4 | 0.0 | 0.03 | 0.23 | 0.24 | 0.23 | 0.17 | 0.17 | -- | -- | -- | -- | -- | -- | -- | -- | -- | 0.05 | -- | 0.05 |
| *G. gallus* | 0.3 | 0.3 | 0.4 | 0.4 | 0.0 | 0.23 | 0.24 | 0.23 | 0.17 | 0.17 | -- | -- | -- | -- | -- | -- | -- | -- | -- | 0.05 | -- | 0.05 |
| *D. melanogaster* | 0.8 | 0.8 | 0.8 | 0.8 | 0.8 | 0.0 | 0.24 | 0.25 | 0.17 | 0.17 | -- | -- | -- | -- | -- | -- | -- | -- | -- | 0.05 | -- | 0.05 |
| *S. cerevisiae* | 1.3 | 1.3 | 1.3 | 1.3 | 1.3 | 1.3 | 0.0 | 0.24 | 0.17 | 0.17 | -- | -- | -- | -- | -- | -- | -- | -- | -- | 0.05 | -- | 0.05 |
| *C. elegans* | 0.8 | 0.8 | 0.8 | 0.8 | 0.8 | 0.7 | 1.3 | 0.0 | 0.17 | 0.17 | -- | -- | -- | -- | -- | -- | -- | -- | -- | 0.05 | -- | 0.05 |
| *A. thaliana* | 1.5 | 1.5 | 1.5 | 1.5 | 1.5 | 1.5 | 1.5 | 1.5 | 0.0 | 0.05 | -- | -- | -- | -- | -- | -- | -- | -- | -- | 0.05 | -- | 0.05 |
| *O. sativa* | 1.5 | 1.5 | 1.5 | 1.5 | 1.5 | 1.5 | 1.5 | 1.5 | 0.2 | 0.0 | -- | -- | -- | -- | -- | -- | -- | -- | -- | 0.05 | -- | 0.05 |
| *B. subtilis* | 4.0 | 4.0 | 4.0 | 4.0 | 4.0 | 4.0 | 4.0 | 4.0 | 4.0 | 4.0 | 0.0 | 0.10 | 0.14 | 0.08 | 0.14 | 0.14 | 0.14 | 0.14 | 0.14 | -- | -- | -- |
| *T. maritima* | 4.0 | 4.0 | 4.0 | 4.0 | 4.0 | 4.0 | 4.0 | 4.0 | 4.0 | 4.0 | 4.0 | 0.0 | 0.10 | 0.10 | 0.10 | 0.10 | 0.10 | 0.10 | 0.10 | -- | -- | -- |
| *P. aeruginosa* | 4.0 | 4.0 | 4.0 | 4.0 | 4.0 | 4.0 | 4.0 | 4.0 | 4.0 | 4.0 | 3.1 | 4.0 | 0.0 | 0.14 | 0.08 | 0.08 | 0.08 | 0.08 | 0.07 | -- | -- | -- |
| *M. tuberculosis* | 4.0 | 4.0 | 4.0 | 4.0 | 4.0 | 4.0 | 4.0 | 4.0 | 4.0 | 4.0 | 3.1 | 4.0 | 3.1 | 0.0 | 0.14 | 0.14 | 0.14 | 0.14 | 0.14 | -- | -- | -- |
| *Y. pestis* | 4.0 | 4.0 | 4.0 | 4.0 | 4.0 | 4.0 | 4.0 | 4.0 | 4.0 | 4.0 | 3.1 | 4.0 | 1.4 | 3.1 | 0.0 | 0.15 | 0.00 | 0.00 | 0.07 | -- | -- | -- |
| *H. influenzae* | 4.0 | 4.0 | 4.0 | 4.0 | 4.0 | 4.0 | 4.0 | 4.0 | 4.0 | 4.0 | 3.1 | 4.0 | 1.4 | 3.1 | 0.7 | 0.0 | 0.15 | 0.15 | 0.07 | -- | -- | -- |
| *S. enterica* | 4.0 | 4.0 | 4.0 | 4.0 | 4.0 | 4.0 | 4.0 | 4.0 | 4.0 | 4.0 | 3.1 | 4.0 | 1.4 | 3.1 | 0.4 | 0.7 | 0.0 | 0.01 | 0.07 | -- | -- | -- |
| *E. coli* | 4.0 | 4.0 | 4.0 | 4.0 | 4.0 | 4.0 | 4.0 | 4.0 | 4.0 | 4.0 | 3.1 | 4.0 | 1.4 | 3.1 | 0.4 | 0.7 | 0.1 | 0.0 | 0.07 | -- | -- | -- |
| *A. fabrum* | 4.0 | 4.0 | 4.0 | 4.0 | 4.0 | 4.0 | 4.0 | 4.0 | 4.0 | 4.0 | 3.1 | 4.0 | 2.5 | 3.1 | 2.5 | 2.5 | 2.5 | 2.5 | 0.0 | -- | -- | -- |
| *P. horikoshii* | 3.8 | 3.8 | 3.8 | 3.8 | 3.8 | 3.8 | 3.8 | 3.8 | 3.8 | 3.8 | 4.0 | 4.0 | 4.0 | 4.0 | 4.0 | 4.0 | 4.0 | 4.0 | 4.0 | 0.0 | 0.29 | 0.10 |
| *S. solfataricus* | 2.7 | 2.7 | 2.7 | 2.7 | 2.7 | 2.7 | 2.7 | 2.7 | 2.7 | 2.7 | 4.0 | 4.0 | 4.0 | 4.0 | 4.0 | 4.0 | 4.0 | 4.0 | 4.0 | 3.8 | 0.0 | 0.29 |
| *H. volcanii* | 3.8 | 3.8 | 3.8 | 3.8 | 3.8 | 3.8 | 3.8 | 3.8 | 3.8 | 3.8 | 4.0 | 4.0 | 4.0 | 4.0 | 4.0 | 4.0 | 4.0 | 4.0 | 4.0 | 3.7 | 3.8 | 0.0 |

^a^ Divergence times were obtained from literature references in the TimeTree database[17].

^b^ Gray shaded cells above the diagonal indicate the standard deviation (in billions of years) of estimated divergence times across studies.

^c^ Divergence times between bacteria and eukaryotes, and between bacteria and archaea were set at 4 billion years (see Methods).

^d^ Divergence times between *S. solfataricus* and eukaryotes were set as the age of the TACK superphylum[64, 65]
